# A Urinary Metabolomic Signature Associated with Small Intestinal Bacterial Overgrowth in Children with Short Bowel Syndrome

**DOI:** 10.64898/2026.11.04.762217

**Authors:** Wenjing Zong, Shuheng Gan, Christina M. Zarek, Miao He, Reena Alame, Elicenda Tovar, Sandra Yang, Samuel Huo, Lindsey Albenberg, Xiaowei Zhan, Gary D. Wu, Christina Bales, Andrew Y. Koh

## Abstract

Direct sampling of the small intestinal microbiome is challenging. Urinary metabolites may offer a noninvasive approach to studying microbial functional activity. We used clinically available gas chromatography–mass spectrometry to profile urinary organic acids in 94 children with short bowel syndrome, comparing those with clinically defined small intestinal bacterial overgrowth (SIBO; n=21) with those without SIBO (n=73). SIBO-associated metabolites included microbial fermentation products, dietary derivatives, and host metabolic intermediates. Metabolites enriched in SIBO mapped to aromatic amino acid metabolism, including tryptophan and tyrosine pathways, consistent with altered microbial fermentation. Many associations persisted after adjustment for intestinal anatomy, age, antibiotic exposure, and nutritional composition. These findings identified a urinary metabolic profile associated with clinically defined SIBO in children with short bowel syndrome and support further evaluation of urinary metabolomics as a noninvasive approach to studying small intestinal microbial function.

## Introduction

The human small intestinal microbiome represents an under-characterized component of the gut microbiome, yet it plays a central role in digestive physiology^1^. The small intestine facilitates most nutrient digestion and absorption, creating a dynamic and interactive biological niche in which dietary substrates, commensal microbes, and host–microbe interactions play critical roles in shaping biological processes. Microbial metabolic activity within the small intestine has potential to directly influence nutrient processing, mucosal health, and systemic metabolism^1^. Therefore, metabolites generated through host–microbe interactions in the small intestine may be critical for understanding health and disease states related to nutrient processing and metabolism.

Small intestinal bacterial overgrowth (SIBO) is a syndrome in which excessive microbial proliferation in the small intestine is associated with disruptions in nutrient absorption, host metabolism and bloodstream infections^2–5^. Children with short bowel syndrome (SBS), the leading cause of pediatric intestinal failure, are particularly susceptible to SIBO due to altered intestinal anatomy, dysmotility, frequent antibiotic exposure, and dependence on parenteral nutrition (PN)^2,5–9^. Diagnosing and monitoring SIBO remains challenging: diagnosis is based on physician intuition, leading to heterogeneous treatments, including broad spectrum antibiotics, with variable responses. Recently, an international expert panel developed the first Delphi consensus-based clinical criteria for SBS-SIBO, providing a standardized framework for defining the clinical phenotype^10^. However, there remains an urgent need for practical, objective measures of SIBO-associated microbial dysfunction that can complement clinical assessment.

Urinary metabolites can provide a functional readout of microbial activity and disease related to gut dysbiosis in the intestine^11–13^. In adults undergoing stem cell transplants, a urinary tryptophan metabolite tracked intestinal microbial disruption and predicted transplant-related mortality^14,15^. Metabolites generated by microbial fermentation in the small intestinal lumen can be absorbed across the intestinal epithelium into circulation and excreted in urine, providing a non-invasive means of sampling microbial activity at a difficult-to-access site. Specifically, several aromatic urinary metabolites have historically been linked to small bowel disease and bacterial overgrowth in SBS^16–21^. Therefore, urinary metabolites may reflect both SIBO-associated microbial activity and the small intestinal nutritional context.

Here, we analyzed urinary metabolite profiles in a discovery cohort of 94 children with SBS. We applied the recently established consensus-based diagnostic criteria in pediatric SBS-SIBO^10^. We hypothesized that children with SIBO would exhibit a distinct urinary metabolite profile reflecting altered small intestinal microbial activity. By integrating metabolite profiling with pathway enrichment analysis, this study demonstrates that urinary organic acid signatures capture a biologically coherent signal of microbial metabolism that is shaped by diet and intestinal anatomy. This work supports urinary metabolomics as a scalable approach for studying small intestinal microbial function in human populations where direct sampling remains impractical.

## Materials and Methods

### Participants and study design

This is a retrospective, cross-sectional cohort study of pediatric patients from birth to 17 years in the Intestinal Rehabilitation Program at the Children’s Hospital of Philadelphia from 2017 to 2022. This research was minimal risk and was deemed exempt by the institution’s IRB board (IRB #17-014008) and was conducted in accordance with the Declaration of Helsinki. Informed consent was waived by the IRB since it was a retrospective study using existing data. Inclusion criteria included history of a urine organic acid assay result, diagnosis of short bowel syndrome based on prior surgical records and imaging, and history of PN dependence for > 2 months. Patients were excluded if they had intestinal failure without a history of surgical resection of the intestine, or conditions that would result in altered urine organic acid profiles including inborn errors of metabolism and hepatic disorders. Medical history, surgical history, dietary history, and medication information were extracted from the electronic medical record system at the time of the clinical urine organic acid assay results. Patients with SIBO were identified by the consensus criteria^10^.

### Sample collection and processing

This study utilized data from historic urine samples that were collected as part of routine clinical care using GC/MS in the CHOP CLIA-certified laboratory (**Supplemental Methods S1**). Metabolite concentrations were reported as milligram per gram of creatinine. Metabolites were annotated by likely source and metabolic pathway using HMDB, KEGG, and primary literature (**Supplemental Methods S2**).

### Analysis

Missing metabolite levels were imputed with half of the minimal observed value for that metabolite. Levels were log_2_-transformed prior to analysis^22^. For the primary differential abundance analysis, we used linear models implemented in the limma R package to identify urinary metabolites associated with SIBO status, with metabolite abundance as the outcome and SIBO status (SIBO vs non-SIBO) as the primary predictor. *P* values were adjusted for multiple testing using the Benjamini-Hochberg false discovery rate (FDR) method (*p*<0.05 considered statistically significant). Model-estimated log_2_ fold changes were calculated for the contrast SIBO minus non-SIBO. Significant metabolites were subjected to KEGG pathway over-representation analysis using MetaboAnalyst (**Supplemental Methods S2**).

In separate metabolite-specific linear regression analyses, we estimated the associations between SIBO status and selected urinary metabolite abundances after adjustment for clinical covariates. Each covariate was evaluated in a separate additive model that included SIBO status, including age, antibiotic exposure, small bowel length remaining, colon resection status, ileocecal valve presence and nutrition composition. Nutrition composition was classified by percent calories derived from each of the three components: food/food-based formula (Food), polymeric formula (EN), and parenteral nutrition (PN), which together sum to 100%. Nutrition composition was represented using two isometric log-ratio (ILR) coordinates, which were included jointly in the model (**Supplemental Methods S3**).

To assess whether urinary metabolites and clinical characteristics could predict SIBO, we developed a random forest classifier and evaluated it using repeated nested cross-validation (CV). Candidate predictors included 79 of 168 metabolites retained after prevalence filtering and seven clinical variables: age, sex, antibiotic exposure, colon resection status, ileocecal valve presence, and percentages of calories from EN and Food. Performance in the outer test folds was assessed using PR-AUC, ROC-AUC, and complementary classification metrics (**Supplemental Methods S4**).

## Results

### Cohort characteristics

We applied the consensus definition for small intestinal bacterial overgrowth (SIBO) in a cross-sectional historic cohort of 94 children with short bowel syndrome (SBS). Baseline demographic and clinical characteristics of the cohort are summarized in **Table 1**.

**Table 1.** Patient demographics and clinical characteristics between SIBO and non-SIBO groups. Food percentage is defined as percentage of total caloric intake derived from foods or food-based formulas consumed. (1. Median (Q1, Q3); n (%); 2. Wilcoxon rank sum test; 3. Pearson’s chi square test. 4. Fisher’s exact test. #Patients could have multiple races or diagnoses.)

|  | **Non-SIBO (n=73)<sup>1</sup>** | **SIBO (n=21)** | **p-value<sup>2</sup>** |
|---|---|---|---|
| Age (years) | 0.3 (0.2, 0.6) | 3.4 (0.9, 5.5) | <0.001<sup>2</sup> |
| Sex, female (%) | 25 (34%) | 9 (43%) | 0.47<sup>3</sup> |
| Race<sup>#</sup> |  |  |  |
| White | 25 (34%) | 12 (57%) | 0.058<sup>3</sup> |
| Black | 26 (36%) | 2 (9.5%) | 0.021<sup>3</sup> |
| Asian | 7 (9.6%) | 3 (14%) | 0.69<sup>4</sup> |
| Other | 20 (27%) | 7 (33%) | 0.60<sup>3</sup> |
| Diagnosis<sup>#</sup> |  |  |  |
| NEC | 32 (44%) | 9 (43%) | 0.94<sup>3</sup> |
| Atresia | 13 (18%) | 2 (9.5%) | 0.51<sup>4</sup> |
| Gastroschisis | 4 (5.5%) | 2 (9.5%) | 0.61<sup>4</sup> |
| Volvulus | 12 (16%) | 5 (24%) | 0.52<sup>4</sup> |
| Hirschsprung | 0 (0%) | 1 (4.8%) | 0.22<sup>4</sup> |
| Others | 27 (37%) | 6 (29%) | 0.48<sup>3</sup> |
| Small bowel remaining (cm) | 30 (21,75) | 40 (24,55) | 0.94<sup>2</sup> |
| Colon resected |  |  | <0.001<sup>3</sup> |
| None | 22 (30%) | 2 (9.5%) |  |
| Partial | 13 (18%) | 18 (86%) |  |
| Total | 38 (52%) | 1 (4.8%) |  |
| Ileocecal valve presence (Yes) | 22 (30%) | 5 (24%) | 0.57<sup>3</sup> |
| Antibiotic status (Yes) | 26 (36%) | 9 (43%) | 0.55<sup>3</sup> |
| Food % | 0 (0,0) | 0 (0,100) | <0.001<sup>2</sup> |

Patients in the SIBO group were older than those in the non-SIBO group (median age 3.4 vs 0.3 years, Wilcoxon rank sum test, p value <0.001). Necrotizing enterocolitis (NEC) was the most common underlying diagnosis in both groups, in line with published literature on SBS epidemiology^9^. Presence of the ileocecal valve (ICV), antibiotic exposure, sex distribution, and the median remaining small bowel length at the last surgery were comparable between groups. Enteral nutrition, based on proportional calories from food/food-based formula and polymeric formulas, was higher in the SIBO group, while most of the non-SIBO group remained fully dependent on parenteral nutrition (PN).

Differences were observed in colon anatomy and nutrition composition. Most patients in the SIBO group had partial colon resection (86%), whereas the non-SIBO group more frequently had total colon resection (52%) (**Table 1**, p value <0.001). Because the ileocecal valve (ICV) is often resected at the same time as the colon, we additionally examined combined colon-ICV anatomy in relation to SIBO status. 53.6% of children without ICV but some colon in continuity (–ICV, + colon) had SIBO, compared to 18.5% of those with both ICV and colon intact, and 2.6% of those missing both ICV and colon (Fischer’s exact test *p*<0.001) (**Supplemental Table 1**). These findings suggest that SIBO risk may depend on the combined intestinal anatomy rather than ICV status alone.

### Identification of differential urinary metabolites associated with SIBO

Prior case series reported several altered urinary metabolites in small intestinal disease and SIBO, suggesting that urine may capture changes in microbial fermentation^16–19^. To determine whether SIBO was associated with a urinary metabolic signature, we performed differential abundance analysis of routine care GC/MS urine organic acid results between SIBO and non-SIBO patients^23,24^. We chose this method for its translational potential, as urinary organic acid assays are widely accessible in many pediatric hospitals and commercial laboratories.

Urinary metabolite profiles demonstrated substantial heterogeneity at the individual level (**Supplemental Figure 1**). 13 urinary metabolites were significantly associated with SIBO status (**Figure 1**, linear model for microarray data, LIMMA. Benjamini-Hochberg FDR *p*-value <0.05), with 6 decreased and 7 increased in SIBO. Additionally, 16 metabolites associated with SIBO status at an unadjusted p-value < 0.05 but did not meet FDR significance (**Supplemental Table 2**).

**Figure 1.**
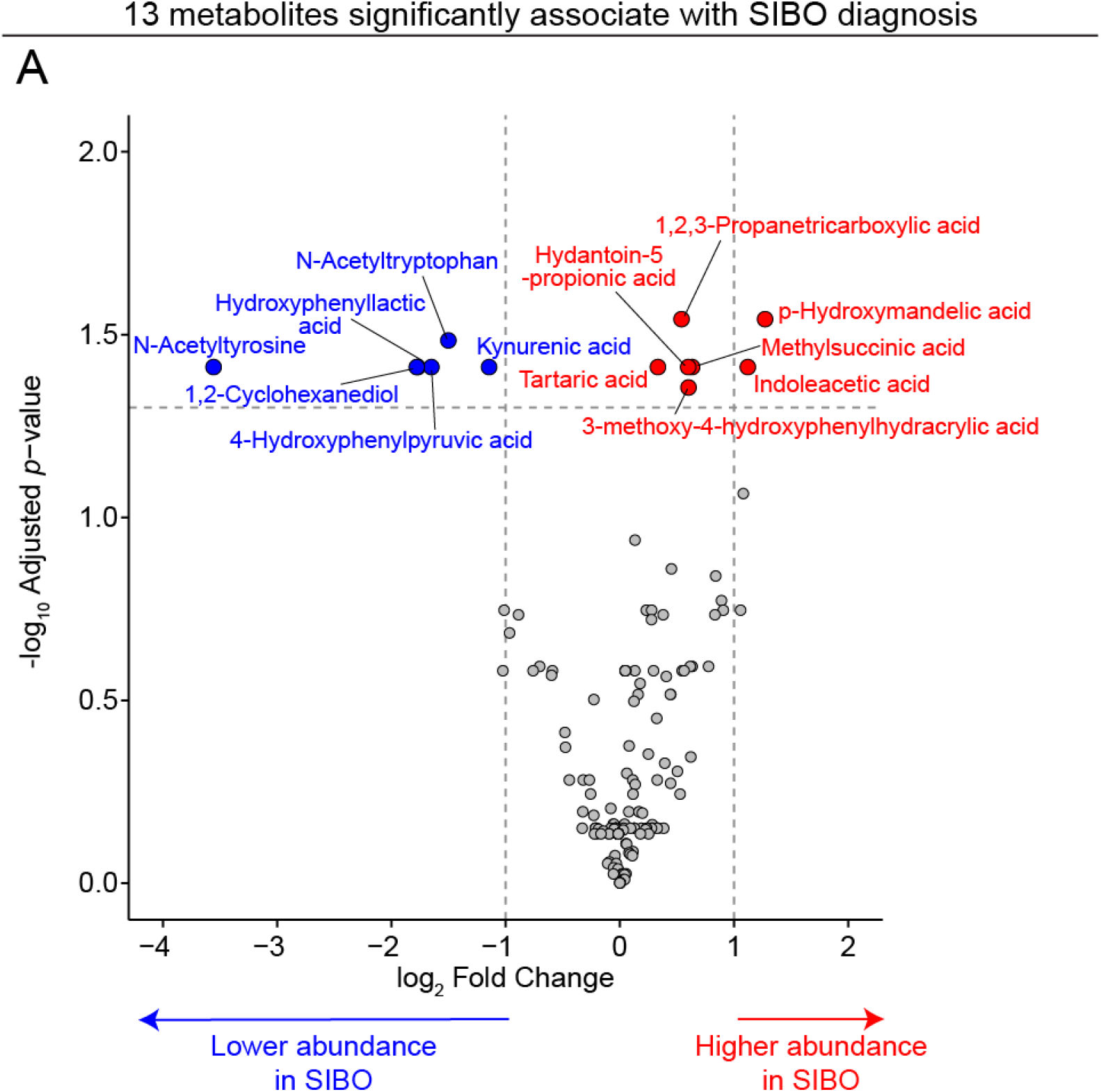
GC-MS analysis of urinary metabolites identifies a signature associated with SIBO. (A) Volcano plot of differential metabolites comparing the SIBO and non-SIBO groups. Metabolites with FDR adjusted *p*-value <0.05 are labeled with metabolite names. Red dots = metabolites with significantly higher abundance in SIBO; blue dots = metabolites with significantly lower abundance in SIBO; gray dots = non-significant metabolites. Dashed, horizontal line = FDR adjusted *p*-value <0.05.

Differential metabolites were broadly categorized into microbial metabolism products (microbial), host metabolic intermediates (host), dietary derivatives (food) or environmental/exposome-associated compounds (exposome) (**Table 2**). Notably, nearly half of the metabolites altered in SIBO were associated with microbial aromatic amino acid metabolism, including the tryptophan and tyrosine fermentation derivatives 4-hydroxyphenyllactic acid (4-HPLA), 4-hydroxyphenylpyruvic acid (4-HPPA), and indoleacetic acid (IAA). Additional metabolites were linked to dietary derivatives and organic acid fermentation from food, such as tartaric acid, suggesting dietary intake increased microbial metabolic activity within the small intestine in SIBO patients. In contrast, several metabolites that were decreased in the SIBO group, including N-acetyl tyrosine and 1,2-cyclohexanediol, were linked to host metabolism, PN components, or medical device related exposures. This pattern is consistent with the lower PN exposure and higher EN and food intake observed in the SIBO group, representing differences in nutrition support and medical device exposure between the groups.

**Table 2.**
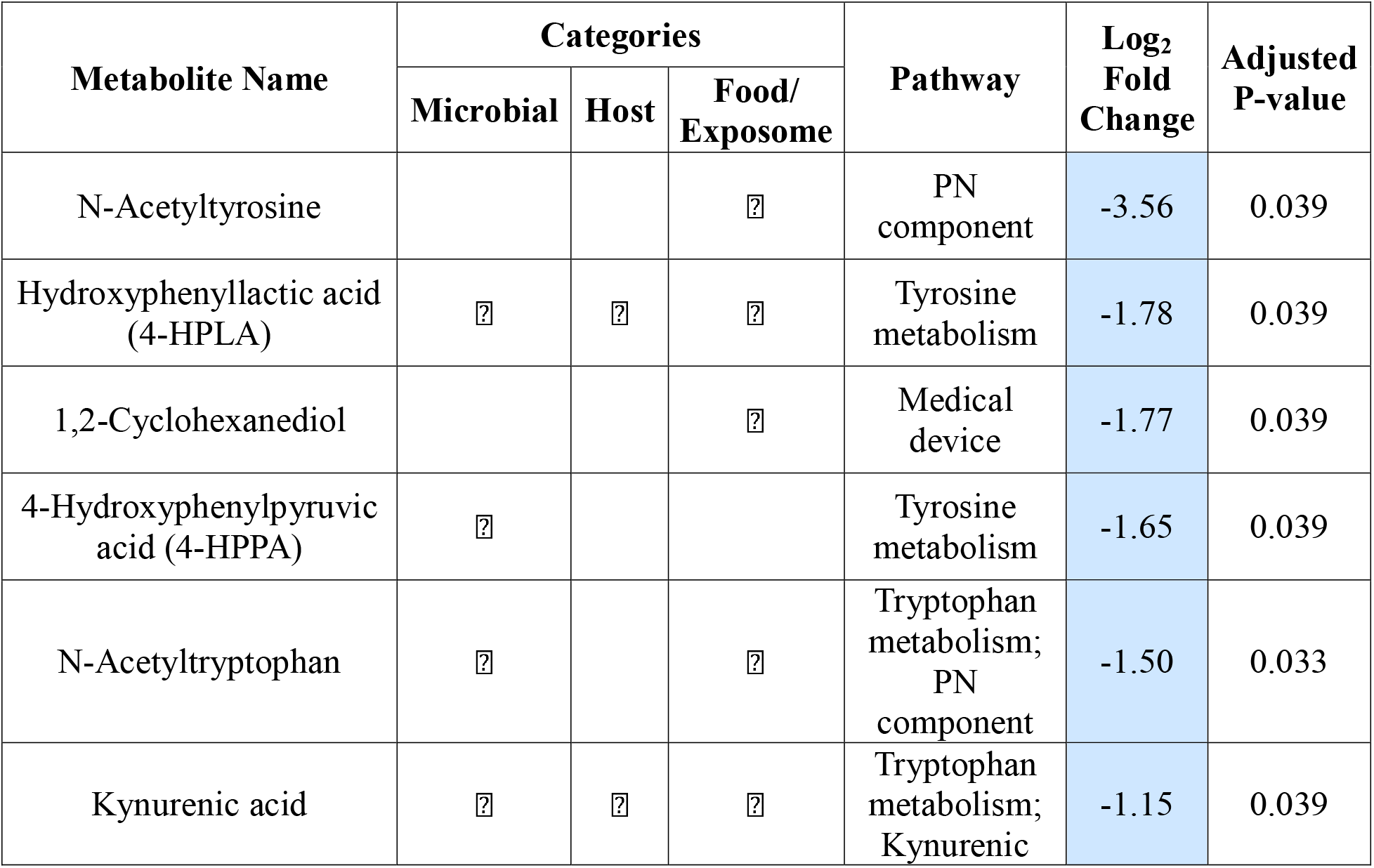
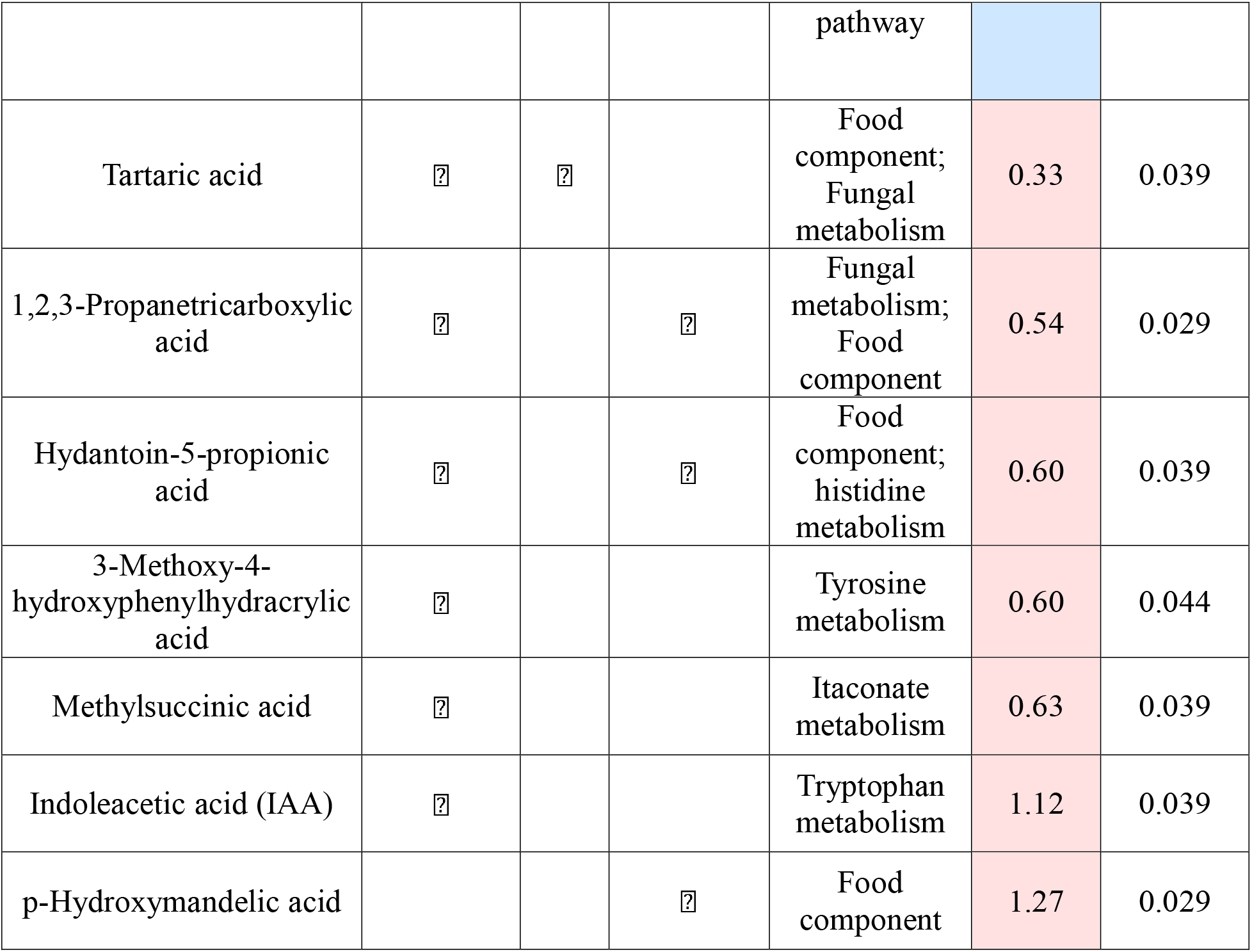
Differential metabolites in SIBO (adjusted *p*<0.05), their assigned categories, pathways, log_2_ fold change (blue=lower, red=higher), and adjusted *p*-values based on FDR correction.

| Metabolite Name | Categories |  |  | Pathway | Log<sub>2</sub> Fold Change | Adjusted P-value |
|---|---|---|---|---|---|---|
|  | Microbial | Host | Food/Exposome |  |  |  |
| N-Acetyltyrosine |  |  | ✓ | PN component | -3.56 | 0.039 |
| Hydroxyphenyllactic acid (4-HPLA) | ✓ | ✓ | ✓ | Tyrosine metabolism | -1.78 | 0.039 |
| 1,2-Cyclohexanediol |  |  | ✓ | Medical device | -1.77 | 0.039 |
| 4-Hydroxyphenylpyruvic acid (4-HPPA) | ✓ |  |  | Tyrosine metabolism | -1.65 | 0.039 |
| N-Acetyltryptophan | ✓ |  | ✓ | Tryptophan metabolism; PN component | -1.50 | 0.033 |
| Kynurenic acid | ✓ | ✓ | ✓ | Tryptophan metabolism; Kynurenic pathway | -1.15 | 0.039 |
| Tartaric acid | ✓ | ✓ |  | Food component; Fungal metabolism | 0.33 | 0.039 |
| 1,2,3-Propanetricarboxylic acid | ✓ |  | ✓ | Fungal metabolism; Food component | 0.54 | 0.029 |
| Hydantoin-5-propionic acid | ✓ |  | ✓ | Food component; histidine metabolism | 0.60 | 0.039 |
| 3-Methoxy-4-hydroxyphenylhydracrylic acid | ✓ |  |  | Tyrosine metabolism | 0.60 | 0.044 |
| Methylsuccinic acid | ✓ |  |  | Itaconate metabolism | 0.63 | 0.039 |
| Indoleacetic acid (IAA) | ✓ |  |  | Tryptophan metabolism | 1.12 | 0.039 |
| p-Hydroxymandelic acid |  |  | ✓ | Food component | 1.27 | 0.029 |

### Pathway analysis reveals enrichment of microbial fermentation pathways

We next performed KEGG pathway analysis using integrated human and bacterial metabolism maps to identify coordinated metabolic changes associated with microbial activity. SIBO-associated metabolites mapped to pathways related to microbial antibiotic metabolism, microbial fermentation, and organic acid metabolism (**Figure 2A**, **Table 2, Supplemental Table 3**). Specifically, multiple metabolites mapped to tryptophan metabolism pathways, including microbial catabolic routes leading to indole derivatives. Additional metabolites were linked to tyrosine metabolism and phenolic compound production, pathways known to be mediated by intestinal microbial communities (**Figure 2B**). Together, these findings suggest that the urinary metabolite profile observed in SIBO reflects altered microbial metabolic activity and substrate utilization within the small intestine.

**Figure 2.**
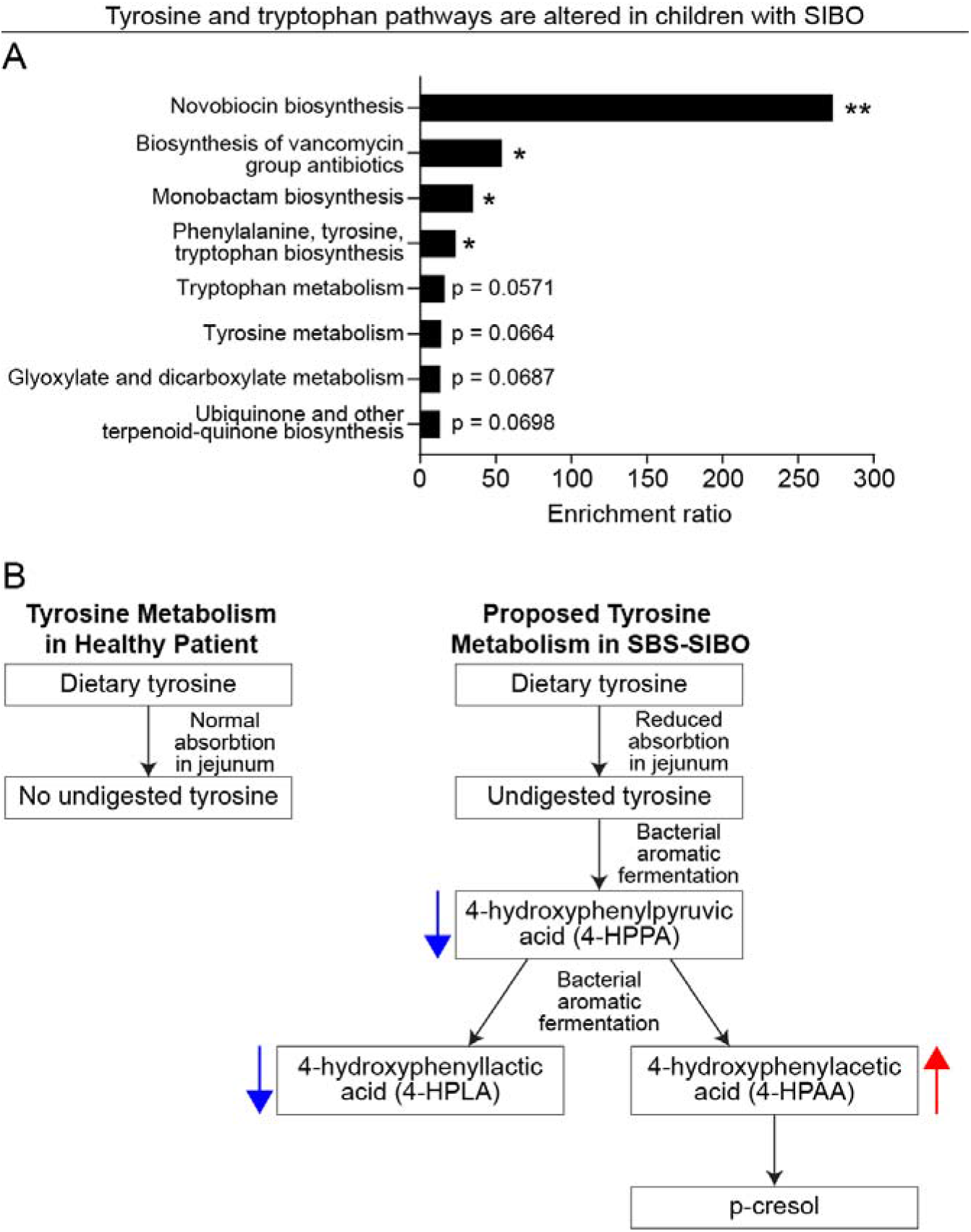
Microbial tyrosine metabolism is enriched in the urine of SBS-SIBO patients. **A)** KEGG pathway enrichment analysis of differentially abundant metabolites. **B)** Tyrosine metabolism in healthy patients and SBS-SIBO patients. Significantly lower (blue arrows) or higher (red arrows) differentially abundant metabolites from our SIBO cohort in the tyrosine metabolic pathway. *Raw *p* < 0.05, **raw *p* < 0.01.

### Assessment of nutrition composition as a clinical covariate

Children with SBS in this cohort received varying combinations of PN, EN (polymeric formula), and food. The host diet is a major contributor to the small bowel luminal environment, where macronutrients and micronutrients impact the structure and metabolic profiles of the small intestinal microbial communities^1^. Furthermore, PN has also been implicated to be strongly associated with dysbiosis and SIBO in SBS children^7^. Thus, nutrition could act either as a confounder or as a biologically relevant modifier that shapes SIBO-associated microbial metabolism^25^. To evaluate the SIBO-associated urinary metabolomic signature after adjusting for nutrition composition, we performed additive linear models using isometric log-ratio (ILR) coordinates for PN, EN, and food as covariate sets to capture degree of PN dependence (ILR1) and composition of enteral intake (ILR2) (**Figure 3**).

**Figure 3.**
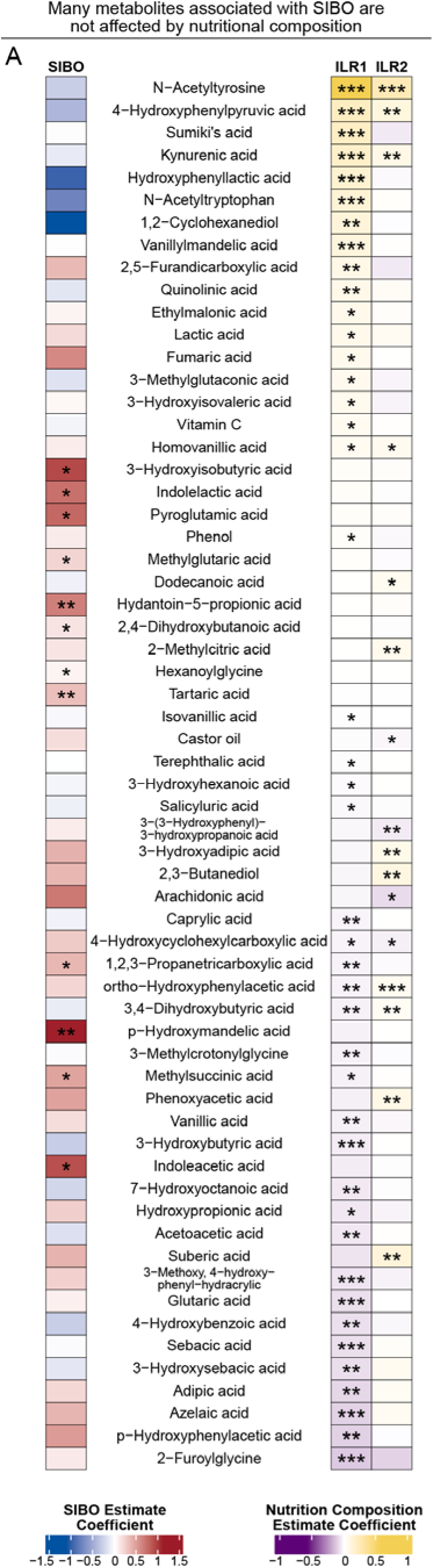
SIBO is associated with a distinct urinary metabolite signature after adjusting for nutrition composition. **A)** Covariate testing of nutritional composition and SIBO status through a linear additive model. Nutrition composition was modeled using isometric log-ratio (ILR) transformation to account for the compositional nature of caloric intake. 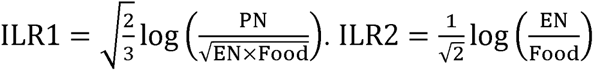 Blue-red scale = SIBO-associated estimate coefficient; strength of SIBO status’s influence on metabolite while accounting for nutritional composition. Purple-yellow scale = nutrition composition estimate coefficient; strength of ILR1’s or ILR2’s influence on metabolite. \**p* < 0.05, \*\**p* < 0.01, \*\*\**p* < 0.001.

Nutrition composition alone significantly impacted a broad range of metabolites (**Figure 3**), including N-acetyltyrosine, N-acetyltryptophan, and 4-HPPA. However, a group of 12 metabolites remained significant for association with SIBO status after nutrition adjustment. Several of these metabolites are microbial fermentation byproducts, including indoleacetic acid, methylsuccinic acid, and hydantoin-5-propionic acid. In summary, nutrition composition impacts many urinary metabolites, though many metabolites remain independently associated with SIBO.

### Assessment of other clinical covariates

We next evaluated age, remaining small bowel length, ileocecal valve presence, colon resection, and antibiotic exposure as clinical covariates in metabolite-specific additive models. In our cohort, children with SIBO were older and more likely to have partial rather than total colon resection (**Table 1**). Age showed a broad metabolomic signal that overlapped with several SIBO-associated metabolites, but the SIBO-associated pattern largely persisted after adjustment (**Figure 4A**). Colon resection was also associated with a subset of metabolites but did not eliminate the SIBO-associated metabolite pattern (**Figure 4B**). In contrast, residual small bowel length and ileocecal valve presence showed fewer independent associations with metabolite abundance (**Figure 4C, 4D**). Antibiotic exposure was similarly distributed between SIBO and non-SIBO groups (**Table 1**) and had minimal association with the metabolite abundance (**Figure 4E**). Together, these analyses suggest that a SIBO-associated urinary metabolite signature persisted after accounting for important clinical covariates, although age and intestinal anatomy contributed to some of the variation in expression of this signature.

**Figure 4.**
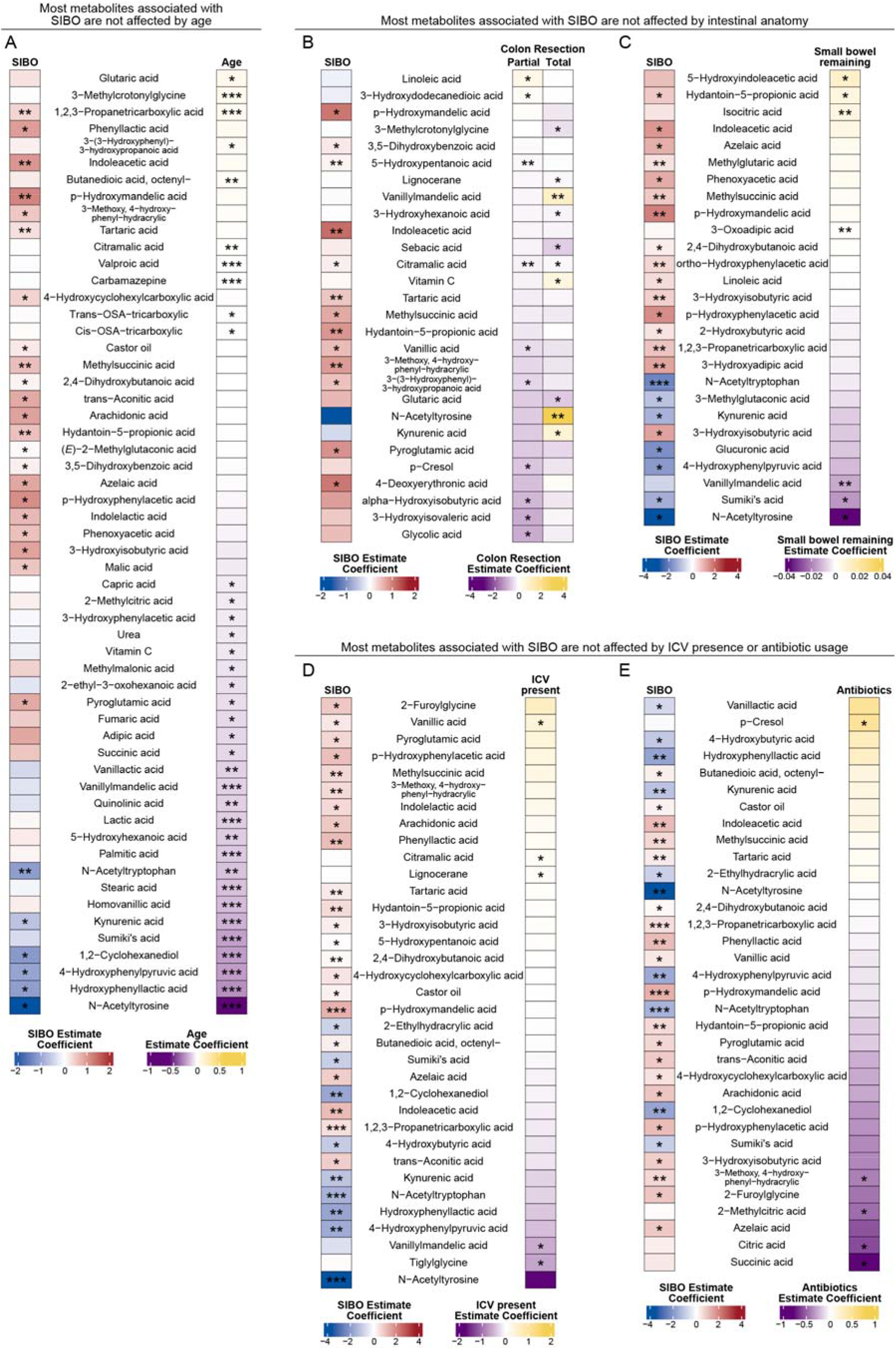
Many of the SIBO-associated metabolites are not affected by clinical covariates, including age, anatomy (SB remaining, ICV status, colon resection) and antibiotics exposure. **A-E)** Covariate testing of clinical variables and SIBO status through a linear additive model. **A)** Age. **B)** Colon resection (total, partial and none). Coefficients shown for partial and total resection represent differences relative to no resection. **C)** Length of remaining small bowel. **D)** ICV presence (yes/no), with ICV present (yes) as the reference category. Significance for ICV present (no) shown. **E)** Antibiotic exposure (yes/no), defined as antibiotic usage within 2 weeks of testing. Significance for antibiotics (yes) shown. Blue-red scale = SIBO-associated estimate coefficient while accounting for each covariate. Purple-yellow scale = covariate estimate coefficient. \**p* < 0.05, \*\**p* < 0.01, \*\*\**p* < 0.001.

### Machine learning corroborates key metabolite and anatomic associations

To evaluate whether the urinary metabolite signature could discriminate SIBO using a multivariable predictive framework, we trained a random forest classifier using 79 candidate urinary metabolites and 7 clinical variables with SMOTE (Synthetic Minority Over-sampling Technique). SMOTE was applied in each training fold to address class imbalance due to fewer SIBO cases (**Figure 5A**). The model discriminated SIBO from non-SIBO better than chance, based on a mean 0.522 Precision-Recall Area Under the Curve (PR-AUC) performance (0.22 baseline for random) (**Figure 5B**). The correct identification of non-SIBO cases, or specificity, was consistently high. In contrast, sensitivity was low (**Figure 5C**). Overall, this model identified SIBO status from a panel of urinary metabolites, supporting the predictive potential of a urinary metabolic signature.

**Figure 5.**
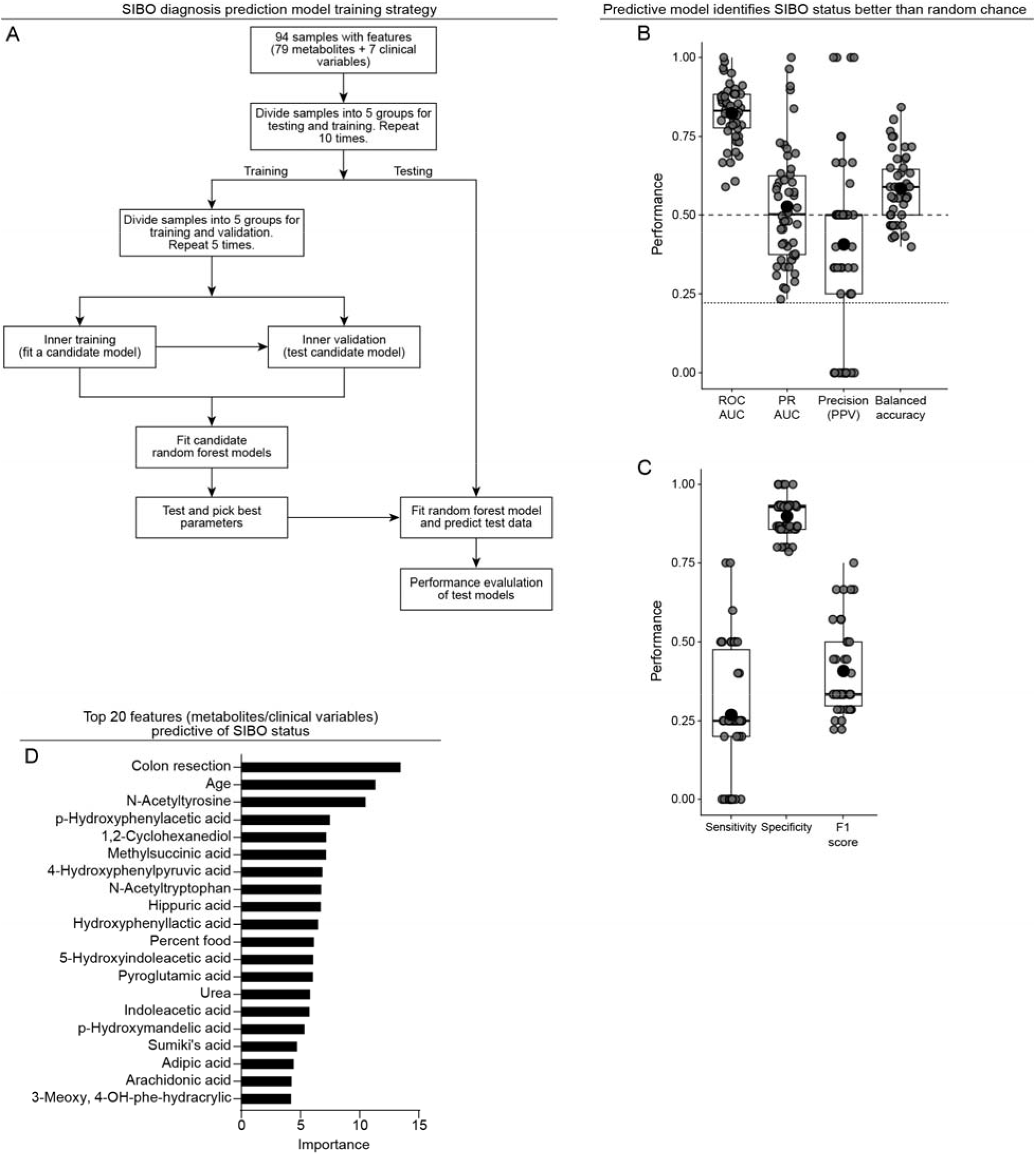
Random forest classification of SIBO using urinary metabolites and clinical variables. **A)** Machine learning training and testing strategy. Features include metabolites and clinical variables. **B)** Performance metrics of the predictive model. Baseline for PR-AUC and precision are 0.22 (21 SIBO samples out of 94 total samples; dotted line). Baseline for ROC-AUC and balanced accuracy are 0.5 (hashed line). Box and whiskers = median, quartiles and extremes. Black dots = mean performance. Grey dots = performance of each run. **C)** Sensitivity, specificity, and F1 score performance of the predictive model. Box and whiskers = median, quartiles and extremes. Black dots = mean performance. Grey dots = performance of each run. **D)** Top 20 features ranked by importance in the random forest model, including urinary metabolites and clinical variables. ROC-AUC, Area Under the Receiver Operating Characteristic Curve; PR-AUC, Precision-Recall Area Under the Curve; Precision (PPV), Precision Positive Predictive Value.

Feature (variable) importance in the model was consistent with two findings from our analyses. Colon resection status was the highest ranked feature among all 86 candidate metabolites and clinical variables, followed by age (**Figure 5D, Supplemental Table 4**). Among the metabolites, tyrosine and tryptophan fermentation metabolites, including 4-hydroxyphenylpyruvic acid (4-HPPA), p-hydroxyphenylacetic acid (4-HPAA), hydroxyphenyllactic acid (4-HPLA), N-acetyltryptophan, and indoleacetic acid (IAA) each ranked among the top 20 features (**Figure 5D**). Overall, this supports our linear model analyses demonstrating that these microbial fermentation metabolites and intestinal anatomy strongly associate with SIBO.

## Discussion

In this study, we identified a distinct urinary metabolite pattern associated with clinically defined SIBO in a discovery cohort of children with short bowel syndrome (SBS). This distinct SIBO-associated signature persisted after accounting for important clinical covariates, including nutritional composition, age, antibiotics, and intestinal anatomy, reflecting the enteral substrate and intestinal anatomy dependent nature of small intestinal microbial metabolism. An independent random forest model corroborated these findings, with colon resection and age as the most important clinical features of SIBO, and tryptophan and tyrosine fermentation metabolites among its highly ranked predictive features.

We found that the SIBO-associated urinary signature was enriched for metabolites involved in microbial tryptophan metabolism. Urinary microbial-host metabolites, particularly tryptophan intermediates, can serve as indicators of gut microbial metabolism and host–microbe metabolic interactions^11–13,16^. Tryptophan-derived kynurenic acid levels in the blood were linked to outcome of PN dependence in a longitudinal pediatric intestinal failure cohort^26^, supporting the broader relevance of aromatic amino acid metabolism to intestinal function. Adult SIBO studies have demonstrated altered urinary tryptophan-pathway metabolites that improve with antimicrobial treatment, supporting the biological relevance of this pathway during bacterial overgrowth^20,21,27^. Additionally, tryptophan-derived indole-3-acetic acid (IAA), enriched in our SIBO cohort, has been shown to act directly on the intestinal epithelium in animal models to upregulate intestinal mucin sulfation and protect against colitis^28^. Microbial tryptophan indoles limit epithelial permeability via a microbiota-dependent mechanism^29^, indicating that intestinal epithelial cells directly sense and respond to microbial tryptophan catabolites^30^. Together, our findings are consistent with altered microbial-host aromatic amino acid metabolism in SBS-SIBO. Whether these metabolites directly influence mucosal barrier function or host-microbe signaling in SBS-SIBO will require future investigation.

The tyrosine and phenylalanine metabolite patterns in SBS-SIBO are similarly supported by prior literature. Pediatric studies linked elevated urinary 4-hydroxyphenylacetic acid to small bowel disease and bacterial overgrowth syndromes, and murine studies demonstrated that urinary aromatic phenolic compounds can originate from gut microbes^16,19^. In prior pediatric studies of SBS and dysbiosis, abnormal urinary 4-hydroxyphenyllactate and phenyllactate were hypothesized to be of bacterial origin^17^. These prior data provide important precedents for our finding that SIBO is associated with a urinary tyrosine and phenolic signature. Additionally, p-cresol, a bacterial tyrosine-fermentation product downstream of p-hydroxyphenylacetic acid (4-HPAA), is a well-recognized marker of SBS-associated microbial dysbiosis^31^. Thus, our finding of elevated 4-HPAA suggests bacterial tyrosine fermentation is increased in SIBO.

Our study also highlights the importance of diet in shaping the urinary metabolome. Children with SBS have highly heterogeneous nutritional compositions, ranging from predominantly PN to increasing proportions of enteral nutrition and food. Diet is a major determinant of gut microbial metabolism, and microbial processing of dietary carbohydrates, polyphenols, and aromatic amino acids leads to a broad range of phenolic and indole metabolites detectable in host biofluids^1,32^. Our results suggest that nutrition composition broadly impacts the metabolic signature, and to some extent, the SIBO signature. Microbial metabolic phenotypes are highly substrate dependent, as the small intestinal microbiome composition and function can be altered by dietary manipulation^33^. This supports a substrate-dependent microbial metabolic framework for SIBO where food is a luminal substrate that impacts the SIBO-associated metabolic signature.

Our findings support that intestinal anatomy may be strongly associated with SIBO-associated metabolite signature. Among the clinical variables examined, colon resection emerged as the most significant structural correlate of the variation in the predictive modeling. This may reflect a scenario of particularly high SIBO risk, in which an available bacterial reservoir in the remaining colon creates potential for reflux into the small intestine without the typical ICV barrier. Further, enteral substrate availability may alter microbial metabolism and create a condition for persistent overgrowth (**Figure 6**). Notably, ICV presence was not significantly associated with the urinary metabolite pattern in our models, in contrast to prior studies showing ICV resection as a SIBO risk factor^7^. This may reflect limited sample size or suggest that the metabolic consequences may depend on the broader anatomic and nutritional context rather than ICV status alone.

**Figure 6.**
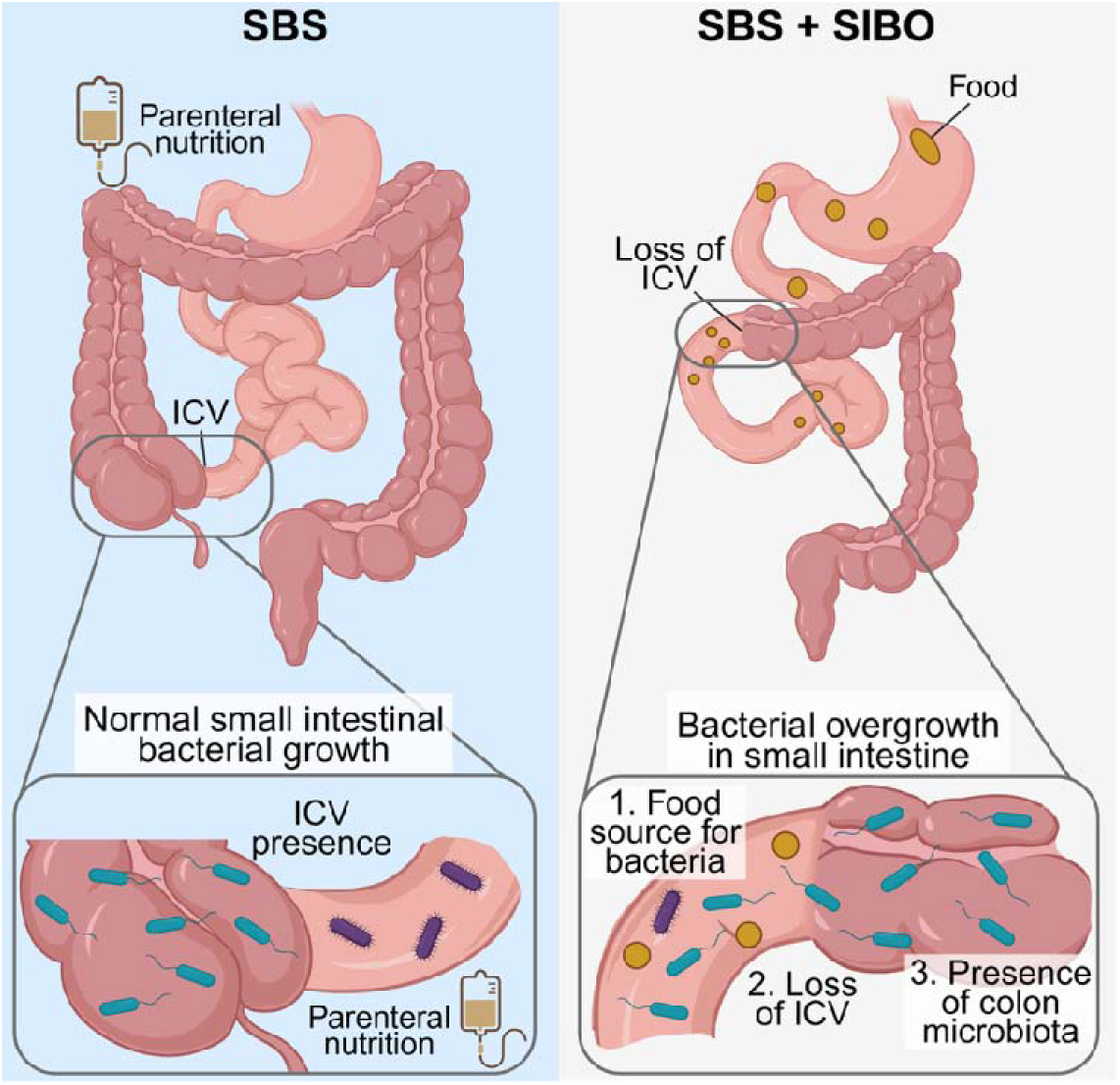
Conceptual framework of nutrition intake, intestinal anatomy and SIBO.

Age contributed broad metabolomic signal in our cohort and was among the top features in the predictive modeling, even though many SIBO-associated metabolites survived adjustment for age. The association between age and metabolites likely reflects more than chronological age alone. In pediatric SBS, age overlaps with developmental stage and therefore feeding advancement and intestinal adaptation. Our exploratory analyses suggested that the relationship between age and nutritional composition differed by SIBO status (data not shown), suggesting that age is not simply a surrogate for diet alone but may function as a composite developmental and clinical context variable. This helps explain that while age impacted a broad range of metabolites, a distinct SIBO-associated signature still remained after adjusting for age.

Our study has several limitations First, this was a single-center retrospective cohort of children with SBS, which may limit generalizability. Second, SIBO status was defined by the recent international consensus rather than a single objective test. This reflects the clinical reality that no objective measure performs well across pediatric SBS populations, but it also introduces the possibility of misclassification. Third, although our analyses explored nutrition composition and several key clinical covariates, residual confounding remains possible. In future studies, it will be important to untangle these covariates through longitudinal sample collections over several SIBO states (active vs resolved) within the same individual. Fourth, the targeted organic acid panel captures only a subset of the urine metabolome. While untargeted approaches may reveal additional pathways not measured in this study, the fact that this targeted platform identified a pathway-coherent signal suggests that it captured biologically meaningful information despite its narrower scope. Fifth, urinary metabolomics were not paired with direct luminal or fecal microbiome sampling. Paired sampling would strengthen mechanistic inference by linking specific metabolites to specific microbial taxa, but direct small bowel luminal sampling was not feasible in this retrospective cohort and is impractical in children. Finally, the random forest model described above should be interpreted as corroborating rather than diagnostic. Lack of external validation and the small number of SIBO cases limit the precision of performance estimates.

These findings have implications for both the study of small intestinal microbial function and the clinical management of SIBO in pediatric SBS. The recent international consensus represents the first effort to standardize SIBO definition in SBS children. Our findings extend this concept by showing that this clinically defined SIBO is associated with a biologically coherent urinary metabolite signature. Urinary metabolomics may offer a noninvasive means of capturing functional microbial activity in the small intestine to complement clinical diagnosis, particularly since stool metagenomics has limited utility in SIBO due to compartment mismatch between colon and small intestine. A key practical strength of this approach is its use of a targeted assay that is already clinically available in commercial labs and pediatric hospitals, offering translational potential without specialized laboratory infrastructure. More broadly, these findings support urinary metabolomics as a scalable strategy for studying small intestinal microbial function in humans where direct sampling remains impractical, a methodologic gap that extends beyond the SBS and SIBO context. Future studies integrating paired luminal and urine profiling, longitudinal sampling across SIBO states, external validation and correlation with clinical outcomes will be essential to establish the stability, specificity, and clinical utility of these metabolite signatures, and to clarify the mechanism through which microbial aromatic amino acid contributes to SIBO pathogenesis and outcomes in SBS.

## Author contributions

W.Z. and C.B. contributed to the conceptualization and design of the study. W.Z., S.G., X.Z., M.H., G.D.W., C.B., and L.A. contributed to methodology, formal analysis, investigation, and editing of the original draft. W.Z. and S.G. performed software development. W.Z. contributed to validation and project administration. W.Z., R.A., E.T., S.Y., and S.H. contributed to data curation and software support. C.M.Z. edited the manuscript and performed visualization. All contributed to critical review of the manuscript, analysis, and validation. G.D.W., C.B., and A.Y.K. contributed to supervision, writing (review and editing), and secured resources. W.Z. is the guarantor of the article. All authors critically revised the manuscript and approved the final version.

## Patient and Public Involvement

Patients and members of the public were not involved in the design, conduct, reporting, or dissemination plans of this research.

## Data Availability Statement

The datasets generated and analyzed from the current study are available from the corresponding author on reasonable request.

## Ethics Statement

This study was reviewed and approved by the Children’s Hospital of Philadelphia Institutional Review Board (IRB #17-014008).

---

**Supplemental Figure 1.**
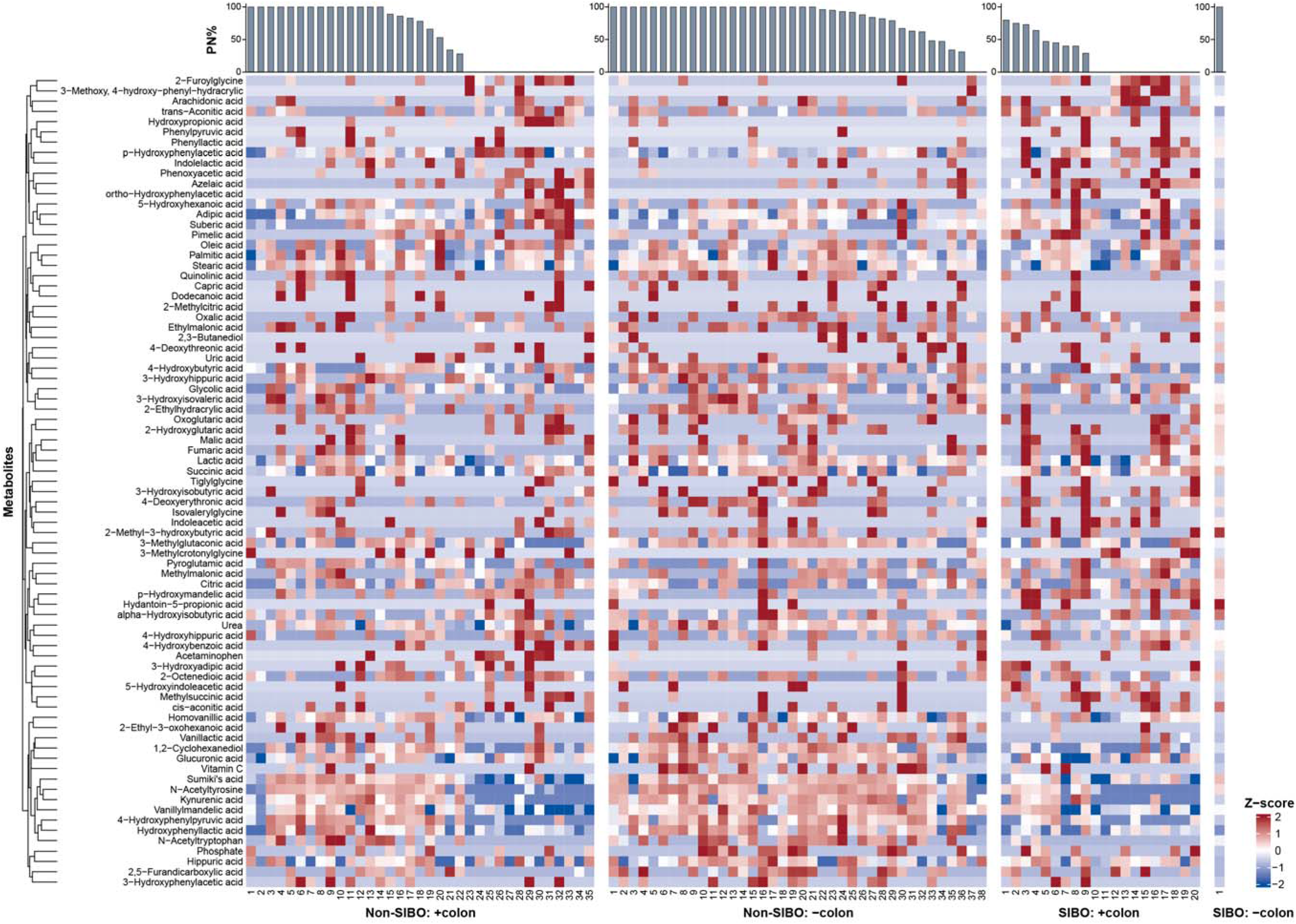
Subject-level urinary metabolite heatmap in children with SBS, with or without SIBO. Profile of urinary metabolites measured by GC/MS across 94 children with SBS (SIBO, n=21; non-SIBO, n=73). Metabolites detected in less than 10 subjects were excluded to reduce sparsity. Rows represent individual metabolites and columns represent individual subjects. Metabolite abundance was displayed as Z-scores, with red indicating higher and blue indicating lower relative abundance. Metabolites were hierarchically clustered and annotated according to microbial, nutrient-related, and other metabolic categories. Subjects were grouped by SIBO status, colon anatomy, and then parenteral nutrition percentage (upper bar graphs).

## Supplemental Tables

**Supplemental Table 1.** Association between intestinal anatomy (ICV presence and colon remaining) and SIBO. Patients were categorized according to ICV presence and colon presence. Colon presence was defined as no or partial colon resection, and colon absence was defined as total colon resection or colon not in continuity. ICV, ileocecal valve.

**Supplemental Table 2.** Differential metabolites in SIBO (adjusted *p* > 0.05, but unadjusted *p* < 0.05) and their assigned categories and pathways.

**Supplemental Table 3.** Metabolite set enrichment analysis of differentially abundant metabolites (LIMMA, adjusted *p* < 0.05). The metabolites were mapped to KEGG pathways from integrated human and gut bacteria metabolism maps. Raw *p*-values were used to determine significance.

**Supplemental Table 4.** Feature importance from the prediction model.

## Supplemental Methods

### S1. Data and GC/MS processing and metabolite identification

Data were collected from the electronic medical record and stored using REDCap, a secure, web-based application. De-identified data was stored on an internal drive.

The urine samples were processed in batches of 10 via Gas Chromatography/Mass Spectrometry (GC/MS), as standard for urinary organic acid panels in the Metabolic Laboratory of the Children’s Hospital of Philadelphia, which is CLIA-certified^34^. Commercially available and National Institute of Standards and Technology (NIST) Mass Spectral Library (Gaithersburg, MD) were utilized to analyze the spectra based on standard operation protocol. Metabolite concentrations were considered acceptable if their quality scores were greater than 80 and their retention times were appropriate. For metabolites with multiple peaks due to different derivatization status, a sum of concentration of peaks with high confidence was taken when appropriate and peaks excluded if the retention time was significantly different from published ranges^34^. This occurred rarely within our samples. Majority of the organic acids were measured semi-quantitatively by comparing the peak area to a single internal standard, 250 mg/g undecanedioic acid. The linearity of presentative organic acids across the 45 min gradient are clinically validated routinely.

### S2. Metabolite annotation and pathway analysis

The metabolic pathway of each metabolite was identified via the Human Metabolite Database (hmdb.ca) when available and cross-checked against published literatures/textbooks^23,34,35^. Metabolites were then annotated into one or more categories based on HMDB and primary literature: microbial, host (e.g., human metabolism intermediary), or food/exposome (e.g., plastic-derived molecules, food metabolites). They were also mapped to KEGG pathways using MetaboAnalyst. KEGG identifiers for stereoisomers were included for metabolites, when possible, as GC/MS cannot distinguish stereoisomers.

Significantly upregulated and downregulated metabolites were analyzed separately using the Over-Representation Analysis (ORA) module for Metabolite Set Enrichment Analysis (MSEA) in MetaboAnalyst (version 6.0) against the KEGG pathway database (November 2025 version).

### S3. Clinical covariate models

Due to the compositional nature of nutritional composition, isometric log-ratio (ILR) transformation was performed using R function “compositions::ilr()” to avoid perfect multicollinearity arising from the constant-sum constraint. Zero values were replaced with 0.01 before ILR transformation.

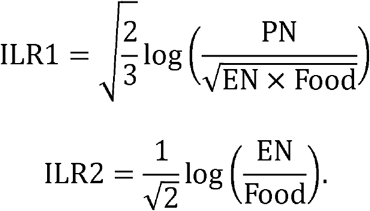

### S4. Random forest

Of the initial panel of 168 urinary metabolites, metabolites detected in fewer than 10 samples were excluded because highly sparse metabolites could be entirely absent from individual training folds. After filtering, 79 metabolites were retained as candidate predictors.

Seven clinical variables were also considered: age, sex, antibiotic exposure, colon resection status, ileocecal valve presence, percentage of calories from enteral nutrition, and percentage of calories from food. Residual small-bowel length was excluded because it was missing for 50 of the 94 participants.

Random forest models were fitted using the R package randomForest version 4.7-1.2. Within each CV split, missing values were imputed using half of the minimum value in the training partition, and the corresponding training-derived value was applied to the held-out partition. Imputation was performed independently within inner and outer CV to prevent data leakage..

Random forest hyperparameters were tuned using 5 times repeated 5-fold inner CV, with the optimal combination selected based on the highest mean precision-recall area under the curve (PR-AUC). Model generalization performance was estimated using 10 times repeated, 5-fold outer CV. Performance measures included PR-AUC, area under the receiver operating characteristic curve (ROC-AUC), sensitivity, specificity, precision, F1 score, and balanced accuracy.

Class imbalance was addressed using the Synthetic Minority Oversampling Technique for mixed continuous and categorical predictors (SMOTENC), implemented using themis::step_smotenc() with over_ratio = 1 and neighbors = 5 (themis version 1.0.3). An over_ratio of 1 increased the minority-class frequency to match that of the majority class. During hyperparameter tuning, SMOTENC was applied exclusively to the training partition of each inner-CV split. After hyperparameter selection, it was applied to the complete outer-training partition before the final model for that outer fold was fitted. The held-out inner-validation and outer-test partitions were not oversampled. Permutation-based feature importance, measured as the mean decrease in prediction accuracy, was extracted from each final outer-fold model using randomForest::importance(). Feature importance values were then averaged across all outer folds and repeats.

